# Nonviral PDMAEMA Vectors Efficiently Express PDE6β in a Mouse Model of Retinitis Pigmentosa

**DOI:** 10.1101/2023.05.04.538567

**Authors:** Diogo B. Bitoque, Sofia M. Calado, Ana M. Rosa da Costa, Gabriela A. Silva

## Abstract

Retinitis Pigmentosa (RP) is one of main causes of inherited blindness, with about 6% of cases caused by a single mutation in the PDE6β gene, making it an ideal candidate for a gene therapy intervention. Gene therapy has been shown to restore normal retinal and visual function in other monogenic diseases, such as Lebers’ Congenital Amaurosis and choroideremia, and RP could benefit from a similar therapeutic approach.

Our goal was to combine efficient nonviral vectors and gene expression systems to express the PDE6β gene in the retina of a mouse model of Retinitis Pigmentosa. We have firstly validated the potential of PDMAEMA polyplexes as a nonviral vector by subretinal injection, which were shown to efficiently promote gene expression in the RPE layer of the adult mouse retina. We have then produced polyplexes of PDMAEMA with a replicating plasmid expressing the PDE6β gene and administered the polyplexes via subretinal injection in the rd10 mouse. We have observed that PDMAEMA polyplexes were able to efficiently enter the target retinal pigment epithelium cells and rapidly express the PDE6β gene in the mouse retina, thus confirming their potential as nonviral vectors for retinal gene therapy.

## 1. Introduction

The aim of gene therapy is to treat diseases by either delivering therapeutic genes to cells or blocking the expression of dysfunctional genes. Viruses have been widely used as vectors for gene therapy due to the natural capacity to infect cells and express their own genome. Despite their high efficiency, viral vectors have limited packaging capacity and most importantly can trigger an immune response that can lead to severe consequences [1]. The drawbacks of viral vectors led to the development non-viral vectors based on lipids and polymers, which are structurally and chemically more versatile, have larger gene packaging capacity and greater stability upon storage and reconstitution [1,2].

Poly(2-(*N,N*-dimethylamino)ethyl methacrylate) (PDMAEMA) is a synthetic polymer whose physicochemical properties have been extensively studied regarding its interaction with DNA [3] and serum proteins [4], its pH-dependent release of nucleic acid load [5], and its long-term stability [6]. We have previously shown that PDMAEMA was capable of complexing genetic material, into nanosized carriers with positive surface charge, and that protect their genetic load from nuclease degradation [7]. PDMAEMA has been shown to efficiently transfect the neural retina [8] and, as we have described, the retinal pigment epithelium [7], which is the ideal target for several retinal pathologies. We have shown that PDMAEMA efficiently enters the cell and traffics through its milieu, reaching the nucleus and releasing its genetic load in the perinuclear space [9].

To test the gene transfer efficiency of PDMAEMA *in vivo*, we have chosen as target a form of Retinitis Pigmentosa (RP) with a mutation in the Rod cGMP-Specific 3,5-Cyclic Phosphodiesterase Beta subunit (PDE6β), which represents about 4-5% of all RP cases [10,11]. This mutation is present in the rd10 mouse, which is considered a model for this form of RP. In this work, we have characterized the phenotypic evolution of RP in this mouse model to determine the most adequate time for administration of the therapeutic vectors. Afterwards we have constructed a PDE6β expression system based on pEPito, an expression plasmid optimized to prevent loss during cell division and less prone to silencing of gene expression [12], which we have previously tested for *in vitro* and *in vivo* gene expression [13,14]. We have combined this plasmid with PDMAEMA to deliver PDE6β to the retina of the rd10 mouse model of RP.

We here present evidence that our PDMAEMA nonviral vector was able to successfully express the PDE6β gene in the retina of the RP mouse model, supporting its potential for gene therapy of the retina.

## 2. Materials and Methods

### 2.1. Plasmid construction

pEPito-eGFP, containing the human CMV enhancer/human elongation factor 1 alpha promoter, was used as backbone [12]. Mouse PDE6β was amplified from pAd.CMVβPDE plasmid (9.9kb, a kind gift from Dr. Jean Bennett, University of Pennsylvania, USA) with specific primers containing NheI (Fw-5’ GATCGCTAGCAATGAGCCTCAGTGAGGAA 3’) and AgeI (Rev-GATCACCGGTTTATAGGATACAGCAGGTCG 3’) restriction sites. The amplified fragment was digested with NheI and AgeI and cloned into pEPito previously digested with the same enzymes. The constructed plasmid (pEPito-PDE6β) with 7.8kb (7814bp, Supplementary figure 1) sequence was confirmed by restriction enzyme digestion and Sanger sequencing, as previously described [15]. pEPito-eGFP (5245bp), encoding the enhanced green fluorescent protein (eGFP) was used as control.

The constructed plasmids were amplified in *E. coli* GT115 competent cells (Invitrogen) in LB medium, extracted and purified using Qiagen’s Maxi-Prep kit. pDNA concentration and quality were determined using a NanoDrop 2000 spectrophotometer (Thermo Scientific) and by agarose gel electrophoresis.

### 2.2. Preparation of PDMAEMA/pDNA polyplexes

A PDMAEMA stock solution of 1 mg/mL was prepared in water and stored at 4°C until further use. Plasmid pEPito-eGFP and pEPito-PDE6β were used to prepare the PDMAEMA:DNA polyplexes. The polyplexes were prepared in water at N/P (nitrogen/phosphorus) ratios of 10:1 and 16:1. These N/P ratios were calculated according to equation 1, where *m*_p_ is the mass of polymer, *m*_D_ is the mass of DNA, *M*_o,D_ is the average repeat unit molecular weight of DNA, and *M*_o,p_ is the repeat unit molecular weight of the polymer.

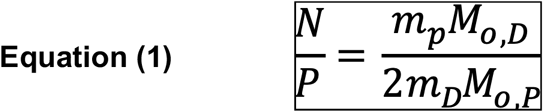

### 2.3. Animals

C57BL/6 (wild-type, WT) and rd10 mice, housed under controlled temperature and a 12-h light/dark cycle with food and water *ad libitum*, were used for the *in vivo* experiments. All experimental procedures were carried out according to the Portuguese and European Union FELASA regulations for the use of animals in research and the Association for Research in Vision and Ophthalmology (ARVO) for the use of animals in ophthalmic and vision research and under the Portuguese national veterinary authority (DGAV) and ethics committee of the host institution. The control group consisted of C57Bl6 animals (n= 3) and the experimental group of rd10 mice (n= 5), harboring the PDE6β mutation [16]. Sample size was calculated (G*Power software) based on [17]. At pre-determined endpoints, animals were humanely sacrificed by administration of anesthesia and subsequent cervical dislocation.

### 2.3 Immunofluorescence of retinal sections

For the characterization of the retinal degeneration, wild-type C57BL/6 and rd10 animals were humanely sacrificed at postnatal days P11, P18, P25 and P35. The eyes were enucleated and fixed in 4% paraformaldehyde (PFA) (Sigma-Aldrich) for 24h, immersed in a gradient of 10%, 20% and 30% sucrose (Sigma-Aldrich) and, after 24 h, included in Optimal Cutting Temperature (OCT) compound (VWR, Radnor, PA, USA). The samples were cryo-sectioned in 10 μm slices using a Leica CM3050 S cryostat (Leica Biosystems) and stored at -20°C.

An immunofluorescence assay was performed in cryosections of rd10 animals using the following antibodies: rabbit polyclonal cone arrestin (1/5000, Merck Millipore), mouse monoclonal rhodopsin (1/1000, Thermo Fisher Scientific) and rabbit polyclonal PDE6β (1/1000, Thermo Fisher Scientific). The retinal sections were washed with 4x PBS/0.25% Triton X-100 (Sigma) and blocked for 1h with 3% goat serum and 1% BSA in PBS/0.25% Triton X-100 (Sigma). The primary antibodies were incubated overnight at 4 °C. After 4x washes with PBS-0.25% Triton X-100 (Sigma), the retinal sections were incubated with secondary antibodies Alexa Fluor® 488 and 594 (Invitrogen) for 1 h at room temperature. After another set of washes, the retinal sections were mounted with Fluoromount G™ containing 2-(4-Amidinophenyl)-1*H*-indole-6-carboxamidine (DAPI, Invitrogen) for nuclei detection, and visualized in a fluorescence (Axio Observer Z2; Zeiss) and confocal microscope (LSM710, Zeiss). Retinal degeneration was quantified by measuring the thickness of the outer nuclear layer (ONL) and of the inner nuclear layer (INL) and calculating the ratio ONL/INL. The transfected area of the retina was calculated by multiplying the length of the region showing transfected cells by the thickness of the retinal slice (10 μm). These quantifications were performed using the Fiji software [18].

### 2.4 Subretinal injection of polyplexes in C57BL/6 and rd10 mice

In order to evaluate the *in vivo* efficiency of the PDMAEMA:DNA polyplexes, 3-month old wild-type C57BL/6 (n=3 per condition) and postnatal day 4 (P4) rd10 mice (n=7 per condition) were injected subretinally in the right eye by using an automatic pump injector (WPI). In WT mice (C57BL/6) 1uL of PDMAEMA polyplexes containing the pEPito-eGFP plasmid, with a N/P ratio of 16:1 (DNA concentration: 0.05 μg/μL), was injected. For rd10 mice a 1 μL solution with each the following conditions was injected: 1 μL of 50 ng/μL and 1 μg/μL concentrations of naked pEPito-hCMV-PDE6β plasmid, injected with and without electroporation; and 1 μL of PDMAEMA polyplexes containing the pEPito-PDE6β plasmid with a N/P ratio of 16:1, with and without electroporation. The left eye was used as control, either non-injected or sham injected. Electroporation was performed using 7 mm tweezer electrodes (Tweezertodes, Havard Apparatus) connected to a BTX ECM 830 (Harvard Apparatus [19,20], under a stereomicroscope (Zeiss, Zeiss Stemi DV4 Stereo Microscope). The electrodes were placed with appropriate orientation for plasmid DNA (negatively charged) and PDMAEMA polyplexes (positively charged) to target the photoreceptor layer after the subretinal injection. The injected animals were humanely sacrificed 14- and 31-days post-injection.

### 2.5 Statistical analysis

The statistical analysis was performed using the Graph Prism 6 software. Statistical analysis for retinal thickness was performed by comparing controls (C57BL/6 mice) and different time-points of the disease model (rd10 mice) using a one-way ANOVA followed by a post hoc Dunnett’s Multiple Comparison test.

## 3. Results and Discussion

### 3.1. PDMAEMA efficiency transfers genes to the mouse retina

Prior to testing a gene of interest *in vivo*, an assessment of the efficiency of the vector should be performed using a reporter gene [21]. To evaluate the efficiency of our vector *in vivo*, we have administered PDMAEMA:pEPito-eGFP polyplexes (P/DNA 16:1) in C57BL/6 mice by subretinal injection. Two weeks after injection, we have observed expression of GFP in cells of the retinal pigment epithelium (RPE) (Figure 1A). GFP expression was further maintained for at least 28 days after injection (Figure 1B). As expected, in the contralateral, non-injected eye we could not observe GPF expression in the retina at 14 (Figure 1C) and 28 days (Figure 1D). The observed transfection profile is similar to our previous study with chitosan that delivered the GFP gene combined with an integrase, which allowed GFP expression for at least 6 months by the RPE [22]. It also shows a similar profile to other synthetic polymers such as PEI transfection of the RPE and neural retina in combination with ultrasound-targeted microbubble destruction, maintained during 21 days after subretinal injection in Sprague-Dawley rats [23].

**Figure 1.** C57BL/6 (WT) mouse eye injected with PDMAEMA polyplexes (P/DNA 16:1) 14 (**A**) and 28 (**B**) days post-injection, and non-injected contralateral eye at 14 (**C**) and 28 (**D**) days. The green color in the injected retinas denotes GFP expression, clearly identifiable when compared with auto-fluorescence from the photoreceptors (box, arrowhead). The retinal detachment observed in both the injected and non-injected eye is a consequence of the process used for cryo-sectioning the retina. N=3. Nuclei stained blue with DAPI. Scale bar: 40 µm.

### 3.2 Retinal degeneration in rd10 mice

To determine the best timeframe for administration of PDMAEMA polyplexes into the retina of rd10 mice, we have evaluated the disease progression in this animal model. We have firstly measured the thickness of the outer nuclear layer (ONL) and of the inner nuclear layer (INL) of the retina. We have observed a significant decrease in retinal thickness after P18 when compared with wild type (WT) animals (Figure 2). Our results are in accordance with other groups that have studied the decrease in the ONL in the animal model, where the decrease of nuclei counts in the ONL, with the same pattern of decrease but at slightly different time-points [24] and through optical coherence tomography (OCT) [11]. We have labeled cone (anti-Cone Arrestin) and rod (anti-Rhodopsin) photoreceptors and observed that the decreased thickness is due to degeneration of the photoreceptor layer after P18. By P35 the outer nuclear layer and the outer segments of cone and rod photoreceptors are almost completely absent (Figure 2). This progressive degeneration phenotype is also in accordance to what is described in the literature [11,25,26].

**Figure 2.**
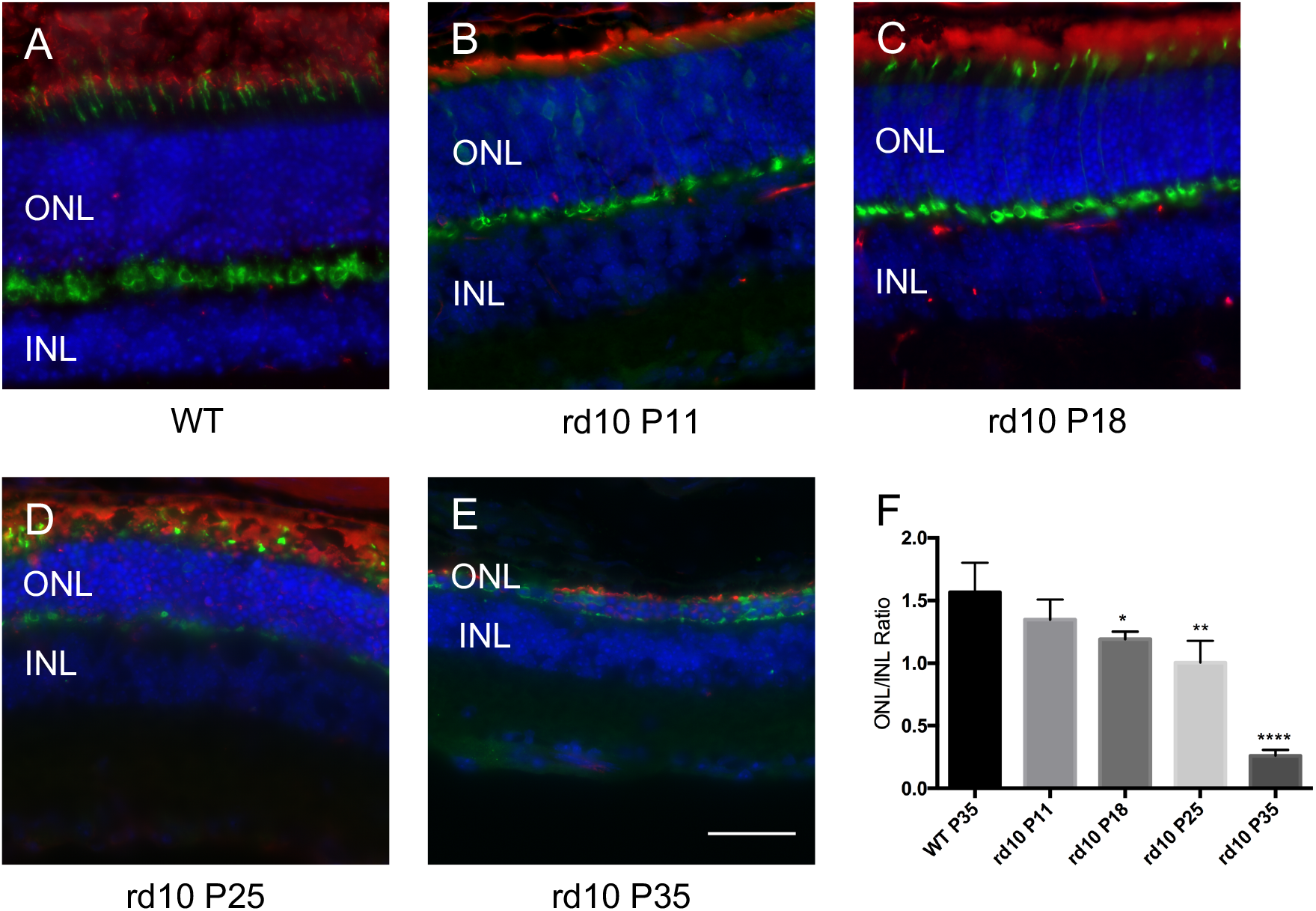
Retinal degeneration of rd10 mice. Immunostaining against rhodopsin as a marker of rods (red), cone-arrestin as a marker of cones (green) and DAPI staining cell nuclei (blue) and structural degeneration of the rd10 retina over time (**B**- P11, **C**- P18, **D**- P25 and **E**- P35) and as control the WT (**A**- P35). N=3. Scale bar: 40 µm; **F**- ONL/INL ratio of P11, P18, P25 and P35 rd10 mice. Statistical significance was determined with one-way ANOVA followed by a post hoc Dunnett’s Multiple Comparison Test to compare the WT mice with the different rd10 timepoints and the statistical difference indicated by the star (*) symbol (* p<0.05; ** p<0.01; *** p<0.001; **** p<0.0001).

### 3.3. PDMAEMA polyplexes deliver PDE6β gene to the retina

Based on the results observed for retinal degeneration, we have set P4 as the time-point to administer the PDMAEMA polyplexes by subretinal injection. The results in Figure 3 show that the lowest concentration of pEPito-PDE6β plasmid (0.05 μg/μL) either alone or in combination with electroporation did not yield detectable gene expression (Figure 3C and 3D). The highest concentration of pEPito-PDE6β plasmid (1 μg/μL) either alone or in combination with electroporation was capable of transfecting retinal cells, as expected (Figure 3E and 3F). The results for naked plasmids are in accordance with what is described in the literature, since it is known that transfection is concentration-dependent and the use of electroporation increases the gene transfer efficiency in the retina [20,27]. Since we have injected a high concentration of free plasmid (1 μg/μL) and the mouse retina only fully matures between P9 and P11, the transfected cells were transfected since they were still dividing at P4 are therefore more permissible to transfection [28]. PDMAEMA polyplexes (N/P ratio 16:1, DNA concentration: 0.05 μg/μL) either alone or in combination with electroporation were also capable of transfecting cells in the rd10 retina (Figure 3G and 3H), but we have not observed differences in the transfection efficiency associated with electroporation. The differences observed between the expression of the pEPito-PDE6β plasmid (0.05 μg/μL) and of PDMAEMA polyplexes can be attributed to PDMAEMA being able to complex and protect the DNA [7]. It is noteworthy that the amount of plasmid used in this study (naked and complexed with PDMAEMA) is significantly less than that compared with other studies [19,20,29], but we still observe a significant number of cells transfected by PDMAEMA polyplexes.

**Figure 3.**
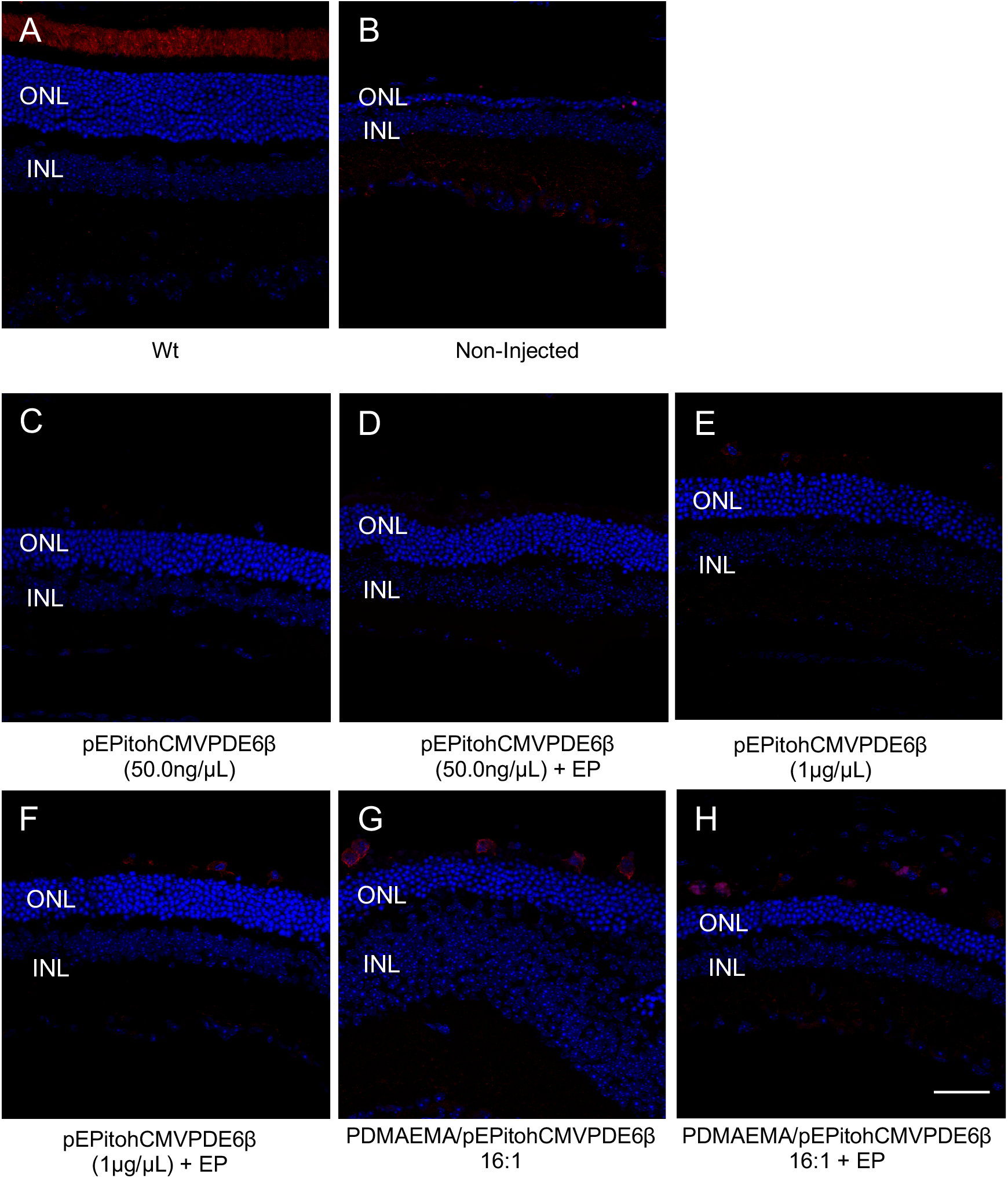
Representative images of C57BL/6 (WT) (**A**), non-injected (**B**) and injected rd10 mice with naked plasmid (0.05 μg/μL and 1μg/μL) and PDMAEMA polyplexes (16:1) with (**D, F, H**) and without (**C, E, G**) electroporation (EP). The cryosections were immunoassayed for PDE6β (red) and nucleus (DAPI, blue). Scale bar= 40 μm.

In this work we have administered our vectors via subretinal injection, known to target the RPE and in some instances, the photoreceptors. To further clarify if the vectors were able to also transfect the latter, we have immunostained the retinal sections for rhodopsin, a marker of rod photoreceptors. We have observed the expression of rhodopsin to be located in the outer segments of rod photoreceptors (Figure 4C) in close proximity with PDE6β expression which indicates that is indeed RPE and not photoreceptors expressing PDE6β. This expression pattern is similar to that observed by Andrieu-Soler *et al* [30], who have injected oligonucleotides (ODNs) by intravitreal injection to correct the point mutation present in the PDE6β gene [30]. We believe that this expression pattern could be translated into a therapeutic effect, since it is known that the intimate interaction between photoreceptors and RPE cells is crucial for photoreceptor survival, as the RPE is responsible for the production of trophic factors (such as cis-retinal), transport of nutrients and ions, and phagocytosis of the outer segments [31,32].

**Figure 4.**
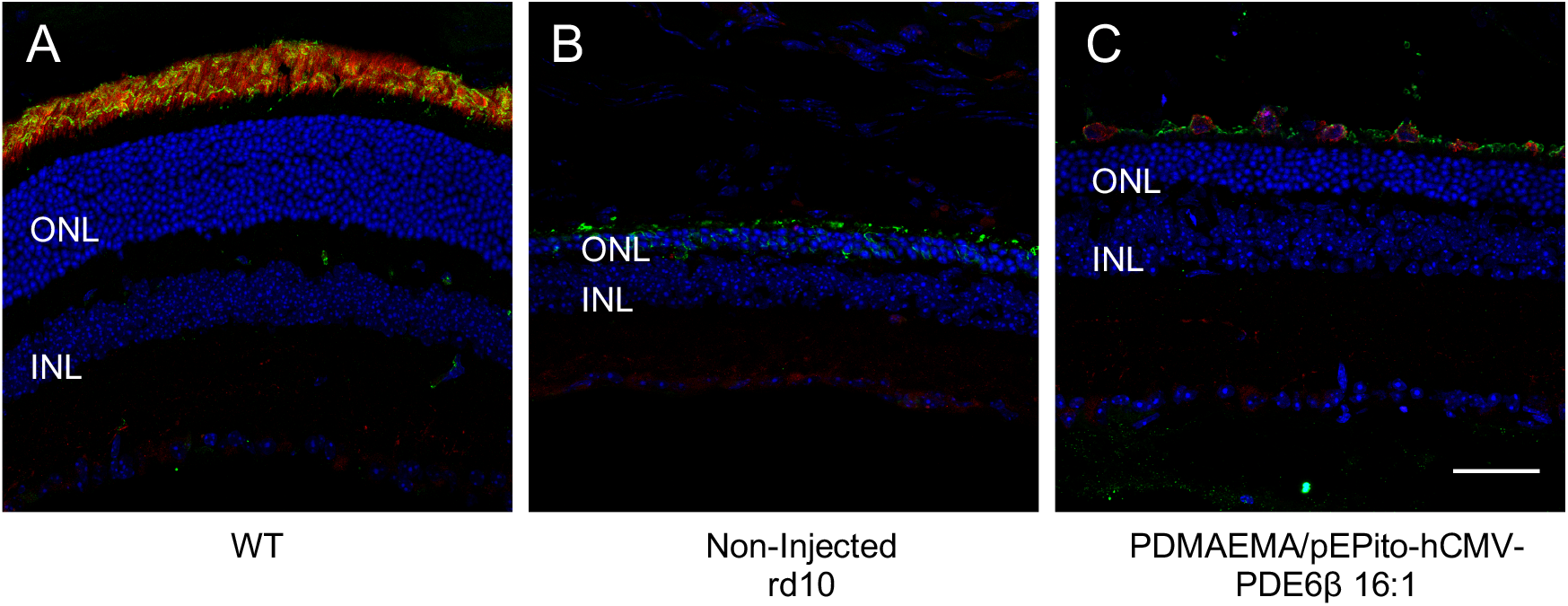
Representative images of C57BL/6 (WT) (**A**), non-injected (**B**) and injected rd10 mice with PDMAEMA polyplexes (16:1) without electroporation (EP) (**C**). Scale bar= 40 μm. The cryosections were immunoassayed for PDE6β (red), Rhodopsin (green) and nucleus (DAPI, blue).

In a photomontage to show a panoramic view of the whole retina (Figure 5) it is possible to observe the wide extension of the PDMAEMA-transfected area. In the literature, it has been described that a transfected area from 0.76mm2 to 8.67mm2 (equivalent to 0.069% to 0.79% of human retina, respectively) in the human eye is sufficient to observe a therapeutic effect [33]. We have measured the transfected area in a cryosection retinal slice corresponding to an area of about 0.00921mm^2^. Considering the total area of the mouse retina [34,35] we were able to estimate the total transfected area of the mouse retina to be 1.54%. This value is consistent to what is sufficient to observe a therapeutic effect.

**Figure 5.**
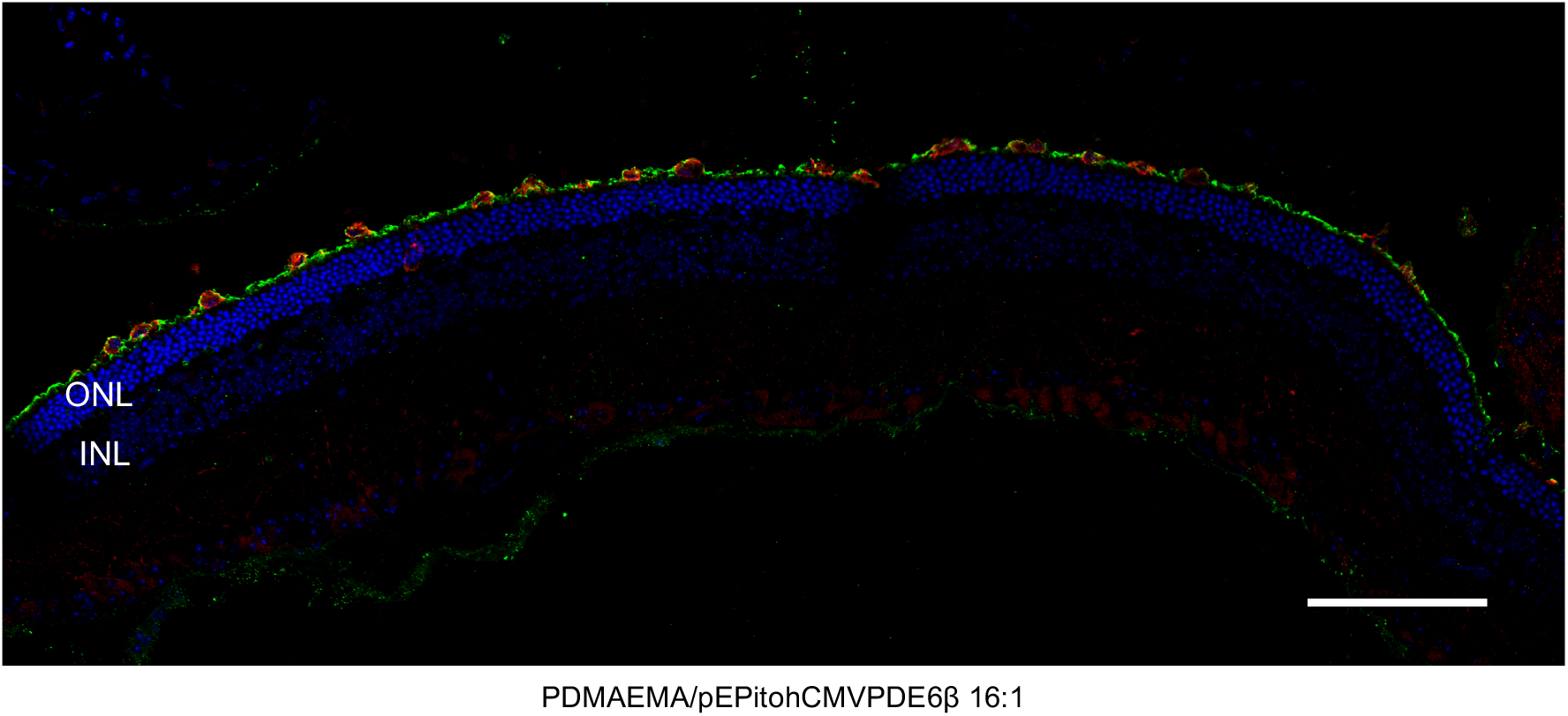
Representative low magnification image of rd10 mice injected with PDMAEMA polyplexes (16:1) without electroporation evidencing several transfected cells. Scale bar= 100 μm. The cryosections were immunoassayed for PDE6β (red), Rhodopsin (green) and nucleus (DAPI, blue).

To further confirm which cells were being transfected by PDMAEMA polyplexes we have also performed immunostaining for the RPE65 protein, a specific marker of RPE cells. As expected, in WT mice the expression of RPE65 does not localize in the same cells as PDE6β (Figure 6A, D and G). In PDMAEMA injected rd10 mice we observed expression of both RPE65 and PDE6β in the same cells (Figure 6C, F and I), which confirms that RPE cells were indeed the ones being transfected by the PDMAEMA polyplexes.

**Figure 6.**
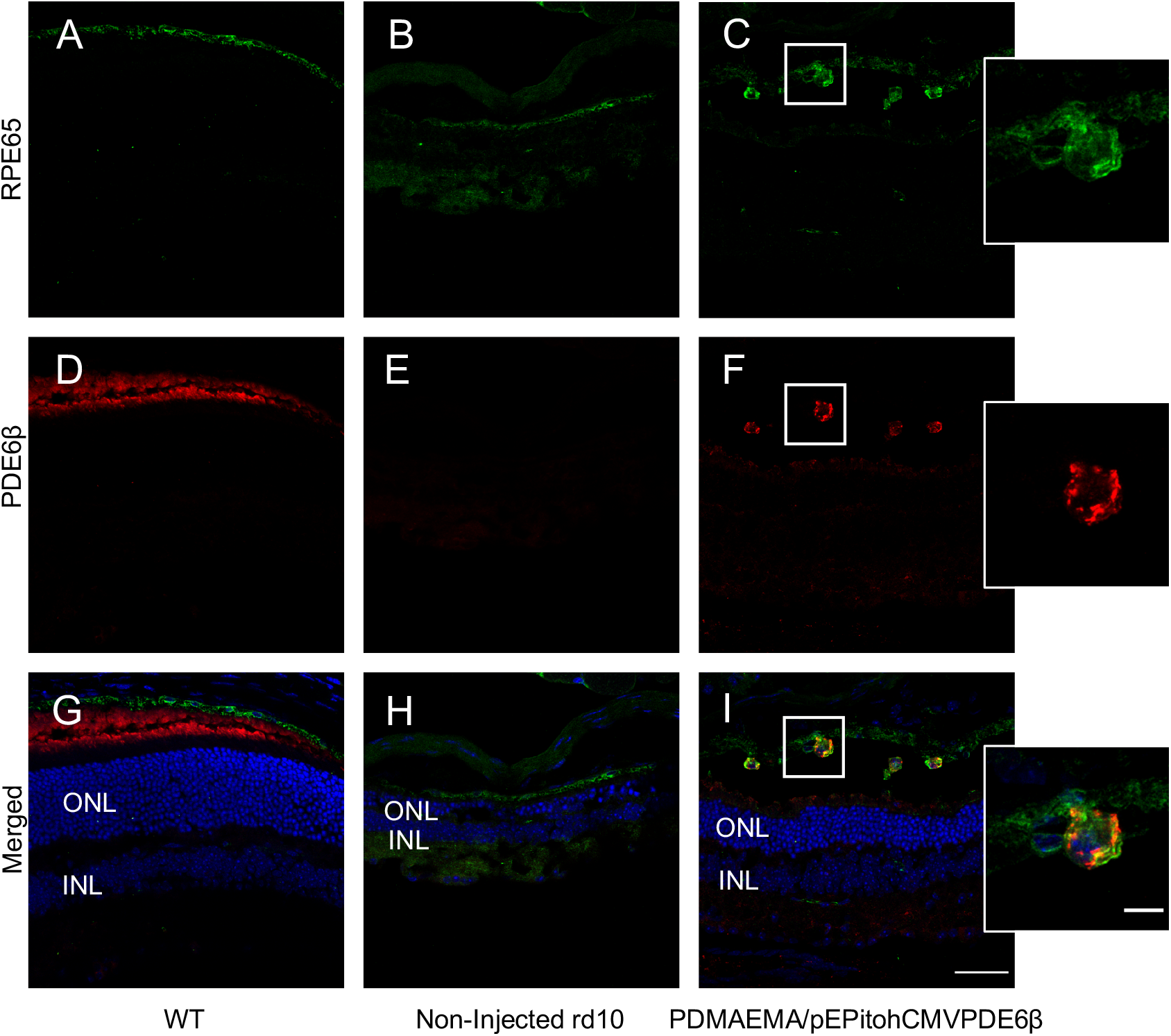
Representative images of C57BL/6 (WT) (**A, D** and **G**), rd10 mice non-injected (**B, E** and **H**) and rd10 mice injected with PDMAEMA polyplexes (16:1) without electroporation (EP) (**C, F** and **I**). Scale bar= 40 μm; Zoom scale bar=10 μm. The cryosections were immunostained for expression of the transfected PDE6β (red), for RPE65 as a RPE marker (green) and the cellular nucleus (DAPI, blue).

Very few gene therapy studies have been performed in rd10 mice, but they have all used viral vectors as delivery agents [36–38], with relative success. As far as we know, we are the first group to report the efficiency of nonviral vectors in delivering the PDE6β gene. This study shows that PDMAEMA is capable of efficiently complex and deliver plasmid DNA to the retina of a mouse model of *retinitis pigmentosa*, and thus constitute an alternative to viral vectors.

## 4. Conclusions

In this study we performed an initial proof of concept study of the efficiency of the non-viral vector Poly(2-(*N,N*-dimethylamino)ethyl methacrylate) (PDMAEMA) polymer in delivering the PDE6β gene to the retina of a mouse model of *retinitis pigmentosa*. After determining the best time for administration, we have shown that PDMAEMA is able to efficiently transfect the RPE in an extensive area, thus pointing to the potential of PDMAEMA polymer as gene vector to the retina. Further studies will focus on assessing the duration of PDE6β expression and in assessing the functional rescue of the retina.

## Acknowledgements

The authors acknowledge the financial support of Fundação para a Ciência e Tecnologia (PTDC/SAU-BEB/098475/2008 and 02/SAICT/2017/028121 to Gabriela A. Silva, PD/BD/52424/2013 individual fellowship to Diogo Bitoque) and the Marie Curie Reintegration Grant (PIRG-GA-2009-249314 to Gabriela A. Silva) under the FP7 program. iNOVA4Health - UID/Multi/04462/2020, a program financially supported by Fundação para a Ciência e Tecnologia / Ministério da Educação e Ciência, through national funds and co-funded by FEDER under the PT2020 Partnership Agreement is also acknowledged.

## References

1. Aied, A.; Greiser, U.; Pandit, A.; Wang, W. Polymer Gene Delivery: Overcoming the Obstacles. Drug Discov Today 2013, 18, 1090–1098, doi:10.1016/j.drudis.2013.06.014.

2. Oliveira, A.V.; Rosa da Costa, A.M.; Silva, G.A. Non-Viral Strategies for Ocular Gene Delivery. Mater Sci Eng C Mater Biol Appl 2017, 77, 1275–1289, doi:10.1016/j.msec.2017.04.068.

3. Arigita, C.; Zuidam, N.J.; Crommelin, D.J.A.; Hennink, W.E. Association and Dissociation Characteristics of Polymer/DNA Complexes Used for Gene Delivery. Pharm Res 1999, 16, 1534–1541, doi:10.1023/A:1015096302720.

4. Cherng, J.-Y.; van de Wetering, P.; Talsma, H.; Crommelin, D.J.A.; Hennink, W.E. Effect of Size and Serum Proteins on Transfection Efficiency of Poly ((2-Dimethylamino)Ethyl Methacrylate)-Plasmid Nanoparticles. Pharm Res 1996, 13, 1038–1042, doi:10.1023/A:1016054623543.

5. Rungsardthong, U.; Ehtezazi, T.; Bailey, L.; Armes, S.P.; Garnett, M.C.; Stolnik, S. Effect of Polymer Ionization on the Interaction with DNA in Nonviral Gene Delivery Systems. Biomacromolecules 2003, 4, 683–690, doi:10.1021/bm025736y.

6. Cherng, J.-Y.; Talsma, H.; Crommelin, D.J.A.; Hennink, W.E. Long Term Stability of Poly((2-Dimethylamino)Ethyl Methacrylate)-Based Gene Delivery Systems. Pharm Res 1999, 16, 1417–1423, doi:10.1023/A:1018907310472.

7. Bitoque, D.B.; Simão, S.; Oliveira, A.V.; Machado, S.; Duran, M.R.; Lopes, E.; Costa, A.M.R. da; Silva, G.A. Efficiency of RAFT-Synthesized PDMAEMA in Gene Transfer to the Retina. Journal of Tissue Engineering and Regenerative Medicine 2017, 11, 265–275, doi:https://doi.org/10.1002/term.1909.

8. Farjo, R.; Skaggs, J.; Quiambao, A.B.; Cooper, M.J.; Naash, M.I. Efficient Non-Viral Ocular Gene Transfer with Compacted DNA Nanoparticles. PLOS ONE 2006, 1, e38, doi:10.1371/journal.pone.0000038.

9. Bitoque, D.B.; Rosa da Costa, A.M.; Silva, G.A. Insights on the Intracellular Trafficking of PDMAEMA Gene Therapy Vectors. Materials Science and Engineering: C 2018, 93, 277–288, doi:10.1016/j.msec.2018.07.071.

10. Francesco, P.; Francesco, S.S.; Diego, P.; Vanessa, B.; Stefano, F.; Enzo, D.I. Retinitis Pigmentosa: Genes and Disease Mechanisms. Current Genomics 2011, 12, 238–249.

11. Pennesi, M.E.; Michaels, K.V.; Magee, S.S.; Maricle, A.; Davin, S.P.; Garg, A.K.; Gale, M.J.; Tu, D.C.; Wen, Y.; Erker, L.R.; et al. Long-Term Characterization of Retinal Degeneration in Rd1 and Rd10 Mice Using Spectral Domain Optical Coherence Tomography. Invest. Ophthalmol. Vis. Sci. 2012, 53, 4644–4656, doi:10.1167/iovs.12-9611.

12. Haase, R.; Argyros, O.; Wong, S.-P.; Harbottle, R.P.; Lipps, H.J.; Ogris, M.; Magnusson, T.; Pinto, M.G.V.; Haas, J.; Baiker, A. PEPito: A Significantly Improved Non-Viral Episomal Expression Vector for Mammalian Cells. BMC Biotechnology 2010, 10, 20, doi:10.1186/1472-6750-10-20.

13. Calado, S.M.; Oliveira, A.V.; Machado, S.; Haase, R.; Silva, G.A. Sustained Gene Expression in the Retina by Improved Episomal Vectors. Tissue Engineering Part A 2014, 20, 2692–2698, doi:10.1089/ten.tea.2013.0672.

14. Calado, S.M.; Diaz-Corrales, F.; Silva, G.A. PEPito-Driven PEDF Expression Ameliorates Diabetic Retinopathy Hallmarks. Human Gene Therapy Methods 2016, 27, 79–86, doi:10.1089/hgtb.2015.169.

15. Bitoque, D.B.; Silva, G.A. Molecular Biology Tools for the Study and Therapy of PDE6β Mutations. Journal of Biotechnology 2018, 284, 1–5, doi:10.1016/j.jbiotec.2018.07.030.

16. 004297 - Pde6b[Rd1-J], Rd10 Strain Details Available online: https://www.jax.org/strain/004297 (accessed on 18 April 2023).

17. Festing, M.F.W.; Altman, D.G. Guidelines for the Design and Statistical Analysis of Experiments Using Laboratory Animals. ILAR Journal 2002, 43, 244–258, doi:10.1093/ilar.43.4.244.

18. Schindelin, J.; Arganda-Carreras, I.; Frise, E.; Kaynig, V.; Longair, M.; Pietzsch, T.; Preibisch, S.; Rueden, C.; Saalfeld, S.; Schmid, B.; et al. Fiji: An Open-Source Platform for Biological-Image Analysis. Nat Methods 2012, 9, 676–682, doi:10.1038/nmeth.2019.

19. Matsuda, T.; Cepko, C.L. Electroporation and RNA Interference in the Rodent Retina in Vivo and in Vitro. Proceedings of the National Academy of Sciences 2004, 101, 16–22, doi:10.1073/pnas.2235688100.

20. Melo, J. de; Blackshaw, S. In Vivo Electroporation of Developing Mouse Retina. JoVE (Journal of Visualized Experiments) 2011, e2847, doi:10.3791/2847.

21. Bennett, J.; Anand, V.; Acland, G.M.; Maguire, A.M. Cross-Species Comparison of in Vivo Reporter Gene Expression after Recombinant Adeno-Associated Virus-Mediated Retinal Transduction. Methods Enzymol 2000, 316, 777–789, doi:10.1016/s0076-6879(00)16762-x.

22. Oliveira, A.V.; Silva, G.A. Chitosan-Based Vectors Mediate Long-Term Gene Expression in the Retina. Journal of Bionanoscience 2015, 9, 373–382, doi:10.1166/jbns.2015.1314.

23. Li, H.; Qian, J.; Yao, C.; Wan, C.; Li, F. Combined Ultrasound-Targeted Microbubble Destruction and Polyethylenimine-Mediated Plasmid DNA Delivery to the Rat Retina: Enhanced Efficiency and Accelerated Expression. The Journal of Gene Medicine 2016, 18, 47–56, doi:10.1002/jgm.2875.

24. Chang, B.; Hawes, N.L.; Pardue, M.T.; German, A.M.; Hurd, R.E.; Davisson, M.T.; Nusinowitz, S.; Rengarajan, K.; Boyd, A.P.; Sidney, S.S.; et al. Two Mouse Retinal Degenerations Caused by Missense Mutations in the β-Subunit of Rod CGMP Phosphodiesterase Gene. Vision Research 2007, 47, 624–633, doi:10.1016/j.visres.2006.11.020.

25. Barhoum, R.; Martínez-Navarrete, G.; Corrochano, S.; Germain, F.; Fernandez-Sanchez, L.; de la Rosa, E.J.; de la Villa, P.; Cuenca, N. Functional and Structural Modifications during Retinal Degeneration in the Rd10 Mouse. Neuroscience 2008, 155, 698–713, doi:10.1016/j.neuroscience.2008.06.042.

26. Gargini, C.; Terzibasi, E.; Mazzoni, F.; Strettoi, E. Retinal organization in the retinal degeneration 10 (rd10) mutant mouse: A morphological and ERG study. Journal of Comparative Neurology 2007, 500, 222–238, doi:10.1002/cne.21144.

27. Nickerson, J.M.; Goodman, P.; Chrenek, M.A.; Bernal, C.J.; Berglin, L.; Redmond, T.M.; Boatright, J.H. Subretinal Delivery and Electroporation in Pigmented and Nonpigmented Adult Mouse Eyes. In Retinal Development: Methods and Protocols; Wang, S.-Z., Ed.; Methods in Molecular Biology; Humana Press: Totowa, NJ, 2012; pp. 53–69 ISBN 978-1-61779-848-1.

28. Young, R.W. Cell differentiation in the retina of the mouse. The Anatomical Record 1985, 212, 199–205, doi:10.1002/ar.1092120215.

29. Johnson, C.J.; Berglin, L.; Chrenek, M.A.; Redmond, T.M.; Boatright, J.H.; Nickerson, J.M. Technical Brief: Subretinal Injection and Electroporation into Adult Mouse Eyes. Mol Vis 2008, 14, 2211–2226.

30. Wang, L.; Xiao, R.; Andres-Mateos, E.; Vandenberghe, L.H. Single Stranded Adeno-Associated Virus Achieves Efficient Gene Transfer to Anterior Segment in the Mouse Eye. PLOS ONE 2017, 12, e0182473, doi:10.1371/journal.pone.0182473.

31. Kevany, B.M.; Palczewski, K. Phagocytosis of Retinal Rod and Cone Photoreceptors. Physiology 2010, 25, 8–15, doi:10.1152/physiol.00038.2009.

32. Strauss, O. The Retinal Pigment Epithelium in Visual Function. Physiol Rev 2005, 85, 845–881, doi:10.1152/physrev.00021.2004.

33. MacLaren, R.E.; Groppe, M.; Barnard, A.R.; Cottriall, C.L.; Tolmachova, T.; Seymour, L.; Clark, K.R.; During, M.J.; Cremers, F.P.M.; Black, G.C.M.; et al. Retinal Gene Therapy in Patients with Choroideremia: Initial Findings from a Phase 1/2 Clinical Trial. The Lancet 2014, 383, 1129–1137, doi:10.1016/S0140-6736(13)62117-0.

34. Remtulla, S.; Hallett, P.E. A Schematic Eye for the Mouse, and Comparisons with the Rat. Vision Res 1985, 25, 21–31, doi:10.1016/0042-6989(85)90076-8.

35. Jeon, C.J.; Strettoi, E.; Masland, R.H. The Major Cell Populations of the Mouse Retina. J Neurosci 1998, 18, 8936–8946, doi:10.1523/JNEUROSCI.18-21-08936.1998.

36. Pang, J.-J.; Boye, S.L.; Kumar, A.; Dinculescu, A.; Deng, W.; Li, J.; Li, Q.; Rani, A.; Foster, T.C.; Chang, B.; et al. AAV-Mediated Gene Therapy for Retinal Degeneration in the Rd10 Mouse Containing a Recessive PDEbeta Mutation. Invest Ophthalmol Vis Sci 2008, 49, 4278–4283, doi:10.1167/iovs.07-1622.

37. Pang, J.; Dai, X.; Boye, S.E.; Barone, I.; Boye, S.L.; Mao, S.; Everhart, D.; Dinculescu, A.; Liu, L.; Umino, Y.; et al. Long-Term Retinal Function and Structure Rescue Using Capsid Mutant AAV8 Vector in the Rd10 Mouse, a Model of Recessive Retinitis Pigmentosa. Mol Ther 2011, 19, 234–242, doi:10.1038/mt.2010.273.

38. Allocca, M.; Manfredi, A.; Iodice, C.; Di Vicino, U.; Auricchio, A. AAV-Mediated Gene Replacement, Either Alone or in Combination with Physical and Pharmacological Agents, Results in Partial and Transient Protection from Photoreceptor Degeneration Associated with ?PDE Deficiency. Investigative Ophthalmology &Visual Science 2011, 52, 5713–5719, doi:10.1167/iovs.10-6269.

